# Microscale Collagen and Fibroblast Interactions Enhance Primary Human Hepatocyte Functions in 3-Dimensional Models

**DOI:** 10.1101/857789

**Authors:** David A. Kukla, Alexandra L. Crampton, David K. Wood, Salman R. Khetani

**Author notes:** Co-corresponding authors **CORRESPONDING AUTHORS’ CONTACT INFORMATION:** Salman R. Khetani, Ph.D., University of Illinois at Chicago, Department of Bioengineering, 851 S Morgan St, 218 SEO, Chicago, IL 60607, David K. Wood, Ph.D., University of Minnesota, Department of Biomedical Engineering, 7-105 Nils Hasselmo Hall, 312 Church St SE, Minneapolis, MN 55455. **Abbreviations:** 2-D: 2-dimensional; 3-D: 3-dimensional; 7-HC: 7-hydroxycoumarin; C_max_: The average maximal concentration the drug reaches in patient’s blood after administration; CEBPa: CCAAT/enhancer binding protein alpha; CYP450: Cytochrome P450 enzymes; DILI: Drug-induced liver injury; DMSO: Dimethylsulfoxide; ECM: Extracellular matrix; ELISA: Enzyme-linked immunosorbent assay; GAPDH: glyceraldehyde 3-phosphate dehydrogenase; HNF4a: hepatocyte nuclear factor 4-alpha; MRP2: multidrug resistance associated protein 2; NPC: Non-parenchymal cell; PEG: Poly(ethylene glycol); PBS: Phosphate buffered saline; PDMS: Polydimethylsiloxane; PHH: Primary human hepatocytes; SLCO1B1: solute carrier organic anion transporter family member 1B1.

## Abstract

Human liver models that are 3-dimensional (3D) in architecture are proving to be indispensable for diverse applications, including compound metabolism and toxicity screening during preclinical drug development, to model human liver diseases for the discovery of novel therapeutics, and for cell-based therapies in the clinic; however, further development of such models is needed to maintain high levels of primary human hepatocyte (PHH) functions for weeks to months *in vitro*. Therefore, here we determined how microscale 3D collagen-I presentation and fibroblast interaction could affect the long-term functions of PHHs. High-throughput droplet microfluidics was utilized to rapidly generate reproducibly-sized (~300 μm diameter) microtissues containing PHHs encapsulated in collagen-I +/− supportive fibroblasts, namely 3T3-J2 murine embryonic fibroblasts or primary human hepatic stellate cells (HSCs); self-assembled spheroids and bulk collagen gels (macrogels) containing PHHs served as gold-standard controls. Hepatic functions (e.g. albumin and cytochrome-P450 or CYP activities) and gene expression were subsequently measured for up to 6 weeks. We found that collagen-based 3D microtissues rescued PHH functions within static multi-well plates at 2- to 30-fold higher levels than self-assembled spheroids or macrogels. Further coating of PHH microtissues with 3T3-J2s led to higher hepatic functions than when the two cell types were either coencapsulated together or when HSCs were used for the coating instead. Additionally, the 3T3-J2-coated PHH microtissues displayed 6+ weeks of relatively stable hepatic gene expression and function at levels similar to freshly thawed PHHs. Lastly, microtissues responded in a clinically-relevant manner to drug-mediated CYP induction or hepatotoxicity. In conclusion, fibroblast-coated collagen microtissues containing PHHs display hepatic functions for 6+ weeks without any fluid perfusion at higher levels than spheroids and macrogels, and such microtissues can be used to assess drug-mediated CYP induction and hepatotoxicity. Ultimately, microtissues may find broader utility for modeling liver diseases and as building blocks for cell-based therapies.

## INTRODUCTION

Owing to significant species-specific differences in drug metabolism functions, *in vitro* models of the human liver are now utilized during preclinical development to assess human-relevant metabolism and toxicity of pharmaceuticals and industrial chemicals ^1^. Such models are also being applied to mimic key cell phenotypes in liver diseases to enable novel drug discovery and to explore the potential of cell-based therapies for patients suffering from end-stage liver disease ^2, 3^. Human liver models can be constructed using transformed cell lines and induced pluripotent stem cell-derived human hepatocytes; however, drug metabolism capacities of both cell sources remain very low ^4^. Additionally, precision cut liver slices lose viability within days due to inflammation ^1^. Therefore, isolated primary human hepatocytes (PHHs) are the ‘gold standard’ for building human liver models ^5^ and novel strategies are being devised to harness *in vivo*-like replicative potential of PHHs towards increasing supply for downstream applications ^6^.

In contrast to rapidly de-differentiating PHHs cultured onto collagen-I adsorbed onto plastic dishes ^7^, several advanced 2-dimensional (2D) and 3-dimensional (3D) culture platforms have been developed to prolong PHH functional lifetime to 2-4 weeks *in vitro*; hepatic functions are typically enhanced in such platforms upon co-culture with liver- or non-liver-derived non-parenchymal cell types ^1^. For instance, when housed in micropatterned co-cultures (MPCCs) with 3T3-J2 murine embryonic fibroblasts, PHHs display a higher level of phenotypic stability than possible with randomly distributed co-cultures of the same two cell types ^7^. However, the adsorbed collagen in MPCCs is insufficient to recapitulate the 3D cell-cell and cell-extracellular matrix (ECM) interactions in physiology and as remodeled in diseases such as fibrosis; furthermore, 2D liver models are not suitable for use as building blocks in regenerative medicine (i.e. cell-based therapies). Similar limitations are apparent in those liver-on-a-chip microfluidic devices that culture PHHs in monolayers on synthetic or natural membranes ^8, 9^. Furthermore, in contrast to static multiwell plates amenable to industrial-scale robotic infrastructure, perfusion-based devices (i.e. liver-on-a-chip) typically reduce the throughput for drug testing, and require fluid handling equipment that adds to assay cost. In contrast, three-dimensional (3D) liver models, such as self-assembled spheroids and bioprinted liver tissues in static devices, can mitigate the above limitations with 2D models. Spheroids, in particular, can be created in different types of multiwell plates for compound screening ^10, 11^. However, it is difficult to form structurally stable spheroids with >50% of PHH donors/lots (and thus spheroid-qualified PHH lots are available at a significant premium from commercial vendors), potentially due to variable ECM secretion rates across donor cells, while bioprinting is an expensive and low-throughput process requiring an unsustainably large number of expensive PHHs for compound screening applications ^12^; therefore, further improvements in 3D human liver models are needed to mitigate the abovementioned limitations.

PHH functions are highly sensitive to the ligation of integrins to proteins within the ECM; thus, encapsulating PHHs within ECM hydrogels at adequate cell densities has been shown to induce liver functions *in vitro* ^5, 13^. However, large (>500 μm) ECM hydrogels that encapsulate cells pose significant diffusion limitations for oxygen and nutrients in the construct’s core in the absence of a functional vasculature ^14^; this limitation can be mitigated by miniaturizing the hydrogel scaffolds to ~100-300 μm, albeit it is not trivial to create reproducible hydrogel scaffolds of this size via manual pipetting. Thus, in recent years, investigators have employed microfluidics to produce highly monodisperse microscale emulsions (droplets) in a high-throughput format ^15, 16^. This so-called ‘droplet microfluidics’ is ideally suited to precisely tune at the microscale cell-cell and cell-ECM interactions and determine optimal conditions for cell survival and function. A few studies have demonstrated the use of droplet microfluidics for liver cell culture. Chen et al encapsulated HepG2 cells in an aqueous droplet surrounded by a crosslinked alginate shell containing NIH-3T3 murine embryonic fibroblasts; the fibroblasts promoted HepG2 albumin and urea secretions as compared to HepG2-only droplets ^17^. Similarly, Siltanen et al developed a coaxial flow-focusing droplet microfluidic device to fabricate microcapsules with a liquid core containing primary rat hepatocytes and a poly(ethylene glycol) (PEG) gel shell; such a droplet configuration promoted tight hepatocyte interactions, which led to albumin/urea secretions and cytochrome-P450 (CYP) 1A1 enzyme activity for 8 days ^18^. Further culturing the hepatocyte droplets atop a 3T3-J2 fibroblast layer led to higher CYP3A activity and albumin secretion for up to 12 days than hepatocyte droplets alone. Finally, Li et al first created 2D micropatterns of primary rat hepatocytes with controlled cell-cell interactions and then lifted such micropatterns using collagenase followed by encapsulation into PEG-based droplets fabricated using a microfluidic device; inclusion of 3T3-J2 fibroblasts into the droplets led to higher albumin secretion and CYP enzyme activities for 16 days ^19^.

While the studies above show the utility of droplet microfluidics for the high-throughput generation of functional hepatocyte/fibroblast droplets using hepatic cell lines and primary rat hepatocytes, the significant differences between these cell types and PHHs in liver functions and drug outcomes necessitates studies with PHHs to determine optimal microenvironmental cues for their long-term culture ^20, 21^. Therefore, here we sought to adapt our previously developed droplet microfluidic device useful for rapidly fabricating protein microgels ^15, 16^ to a) determine the optimal device and culture conditions for the creation of PHH/collagen droplets (microtissues) in a multiwell plate format, b) elucidate the role of different fibroblast populations (3T3-J2 murine embryonic fibroblasts and primary human hepatic stellate cells) on the long-term (up to 6 weeks) enhancement and stabilization of PHH functions when the two cell types were either co-encapsulated within the collagen droplets or when the fibroblasts were ‘coated’ on the outside of the PHH-containing collagen droplets, c) determine the differences in hepatic functional output within microtissues relative to conventional self-assembled spheroids and collagen bulk hydrogels (macrogels), and d) determine the utility of the microtissues for assays in drug development, specifically drug-mediated CYP450 induction and drug-induced hepatotoxicity.

## RESULTS

### Droplet microfluidics for generating reproducible and functional PHH microtissues

PHH microtissues were fabricated using droplet microfluidics and seeded into agarose microwells within industry-standard 24-well plates as illustrated in **Fig. 1a**. While a variety of natural ECM materials are compatible with hepatocyte culture ^5, 13^, rat tail collagen I was selected here for generating microtissues since it is abundantly and cheaply available at high concentrations needed for gelation (>2 mg/mL) and has been utilized extensively for PHH culture without any adverse effects, including the results described below. However, to prevent the aggregation of microtissues before polymerization, microtissues were collected at 37°C directly from the autoclaved microfluidic device (**Fig. 1b**) to ensure immediate polymerization using a bead bath (**Fig 1c,d**). This provided a simple method of polymerizing the microtissues off-chip as opposed to using complicated in-chip heating. The microtissues were collected into an autoclaved 2-mL collection tube, which was directly connected to the microfluidic device with outlet tubing to prevent possible contamination, creating a closed environment between the syringes, microfluidic device, and collection tubes (**Fig. 1d**). The oil was drained from the collection tube using a syringe and microtissues were resuspended in culture medium. Finally, to prevent their aggregation during culture, microtissues were placed into agarose microwells cast within each well of a multi-well plate. Agarose was chosen over PDMS or PEG since it is cheaply available, does not typically interact with compounds of diameters <60nm ^27, 28^, and does not bind to proteins or cells.

**Figure 1.**
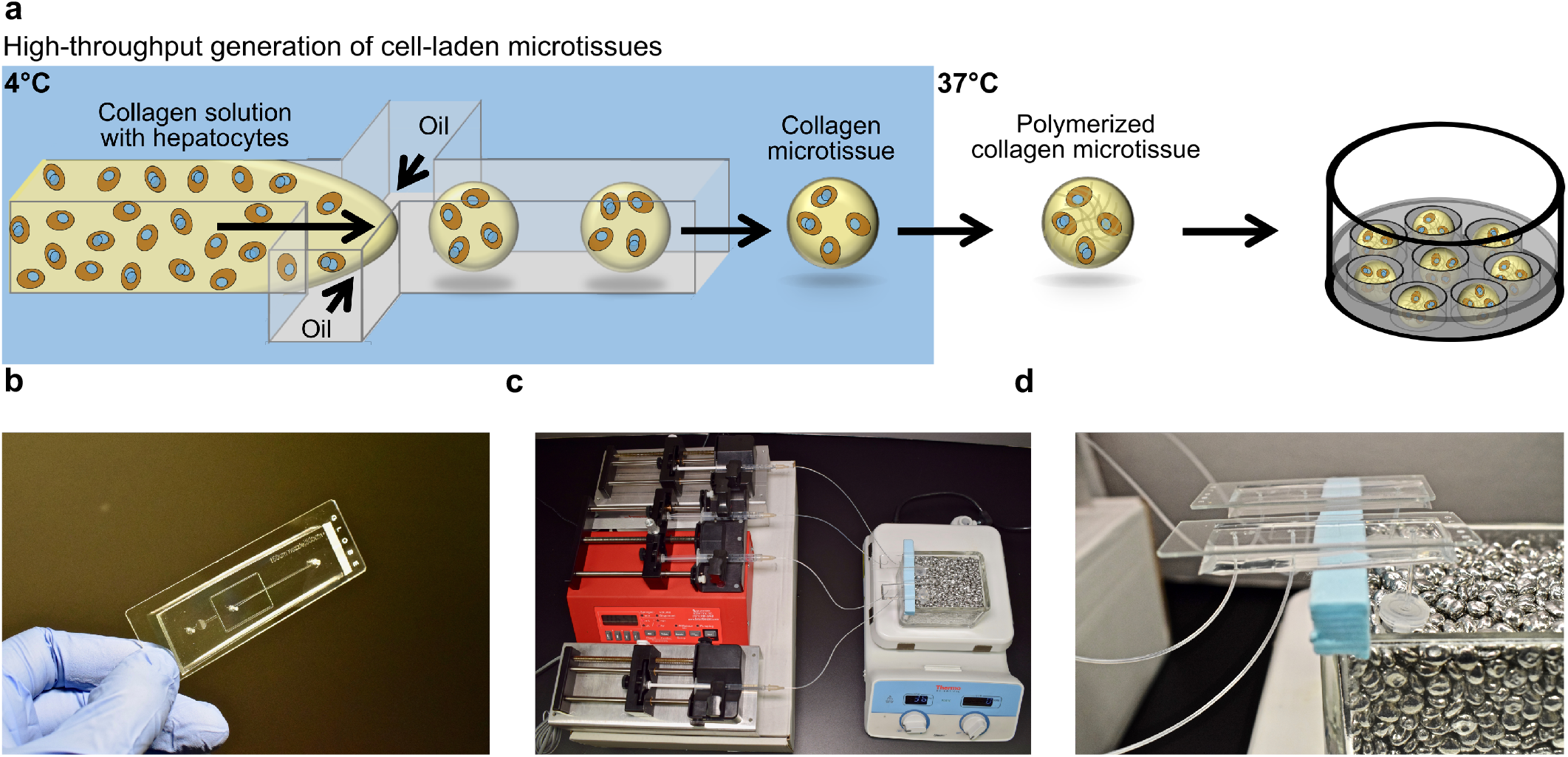
3D human liver tissue platform with tunable cell-cell and cell-ECM interactions for compound screening. **(a)** Hepatocytes are suspended in pH-neutralized collagen solution and then perfused through a droplet generating microfluidic device (see Methods for details about device dimensions). Oil is perfused at a rate ~4 times faster than the aqueous phase to produce microtissues. Microtissues are formed using the microfluidic device at 4°C and collected at 37°C to promote the rapid polymerization of the collagen droplets and encapsulation of the cells within the droplets. Oil is removed, polymerized microtissues are resuspended in culture medium, and subsequently seeded into agarose (2% w/v) microwells cast within multi-well plates. The hepatocytes can be co-cultured with non-parenchymal cell (NPC) types by either co-encapsulating both cell types within the microtissue or by seeding/coating the NPCs onto the surface of the polymerized collagen-based hepatic microtissues. **(b)** Polydimethylsiloxane (PDMS)-based microfluidic devices consisting of a single emulsion droplet generator with 300 μm straight channel and 150 μm nozzle used to fabricate microtissues. **(c)** Microtissue fabrication setup to create microtissues. Two parallel microfluidic devices with syringes containing either oil or collagen+cells are shown; however, additional devices can be run simultaneously to increase fabrication. **(d)** Microfluidic devices with collection tubes. The microtissues are collected into a 2-mL collection tube connected to the microfluidic device with a short outlet tubing. The collection tube is kept at 37°C in a heated bead bath to ensure immediate polymerization of the microtissues as they are dispensed into the tube.

Since PHH functions are known to be dependent on homotypic interactions and resulting tight junction formation in 2D cultures ^7^, here PHHs were mixed with the collagen I at different densities (1.25e6, 2.5e6, 3.75e6, and 5e6 cells/mL) prior to introduction into the droplet microfluidic device. Highly reproducible microtissues were produced with an average diameter of 267.4 μm (standard deviation of 32.7 μm, n=375 microtissues) while containing ~17 and ~34 encapsulated PHHs per microtissue for 1.25e6 (**Fig. 2a**) and 2.5e6 (**Fig. 2b**) PHHs/mL densities, respectively. In contrast, microtissues created using the densities of 3.75e6 or 5e6 PHHs/mL produced several larger microtissues that were not the desired diameter and did not fit into the agarose microwells; furthermore, such high cell densities also caused some instability in the oil emulsion, producing a ‘webbing’ of collagen with entrapped PHHs (data not shown). At a functional level, the microtissues created using 2.5e6 PHHs/mL secreted albumin and displayed CYP3A4 activity at ~2-and 2.5-fold higher rates (normalized to cell number), respectively, than microtissues created using 1.25e6 PHHs/mL after 2 weeks (**Fig. 2c**), thereby showing the positive effects of increased homotypic interactions on hepatic functional output. Thus, the 2.5e6 PHHs/mL density was selected for all subsequent studies with microtissues.

**Figure 2.**
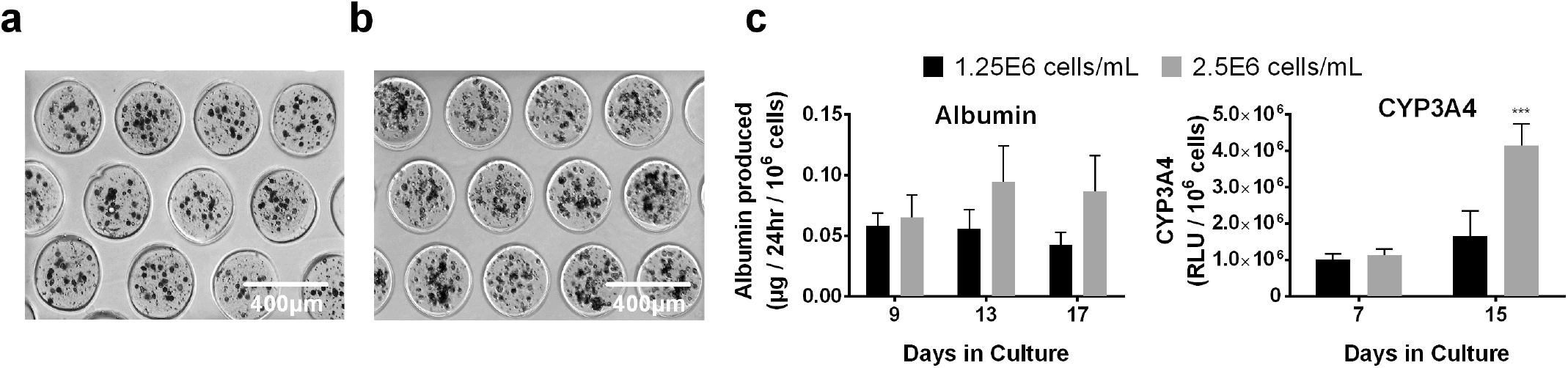
Increasing PHH homotypic interactions within microtissues enhances liver functions. Collagen-based microtissues containing increasing densities of PHHs in a pH-neutralized collagen (1.25e6 and 2.5e6 cells/mL) solution were fabricated using the droplet microfluidic device shown in Fig. 1. Representative phase contrast images of PHH microtissues with a density of **(a)** 1.25e6 cells/mL and **(b)** 2.5e6 cells/mL placed within agarose microwells cast within 24-well plates. Both 1.25e6 and 2.5e6 cells/mL densities produced reproducible microtissues with high yields that fit within the 300 μm x 300 μm agarose microwells. **(c)** Albumin production and CYP3A4 enzyme activity for PHH microtissues fabricated using a cell density of 1.25e6 and 2.5e6 cells/mL. Microtissues with 3.75e6 and 5e6 cells/mL densities were also fabricated, but these densities did not produce homogenously sized microtissues (not shown). Statistical significance is displayed relative to 1.25e6 cells/mL at the same time point. \*\*\**p* ≤ 0.001.

### PHH microtissues functionally outperform self-assembled spheroids and macrogels

Microtissues at 2.5e6 PHHs/mL were created as described above (**Fig. 3a**). Self-assembled spheroids of consistent sizes were created by seeding PHHs directly into the agarose microwells; 200K cells/well (24-well plates) produced tight spheroids with an average diameter of 207.9 μm (standard deviation 21.3 μm, n=375 microtissues) (**Fig. 3b**). Lastly, collagen was dispensed directly into multi-well plates to create macrogels with 20K cells/well, a number of cells that is consistent with that present in the microtissues created at 2.5e6 PHHs/mL (**Fig. 3c**); the collagen hydrogel containing PHHs stably adhered to the plastic plates.

**Figure 3.**
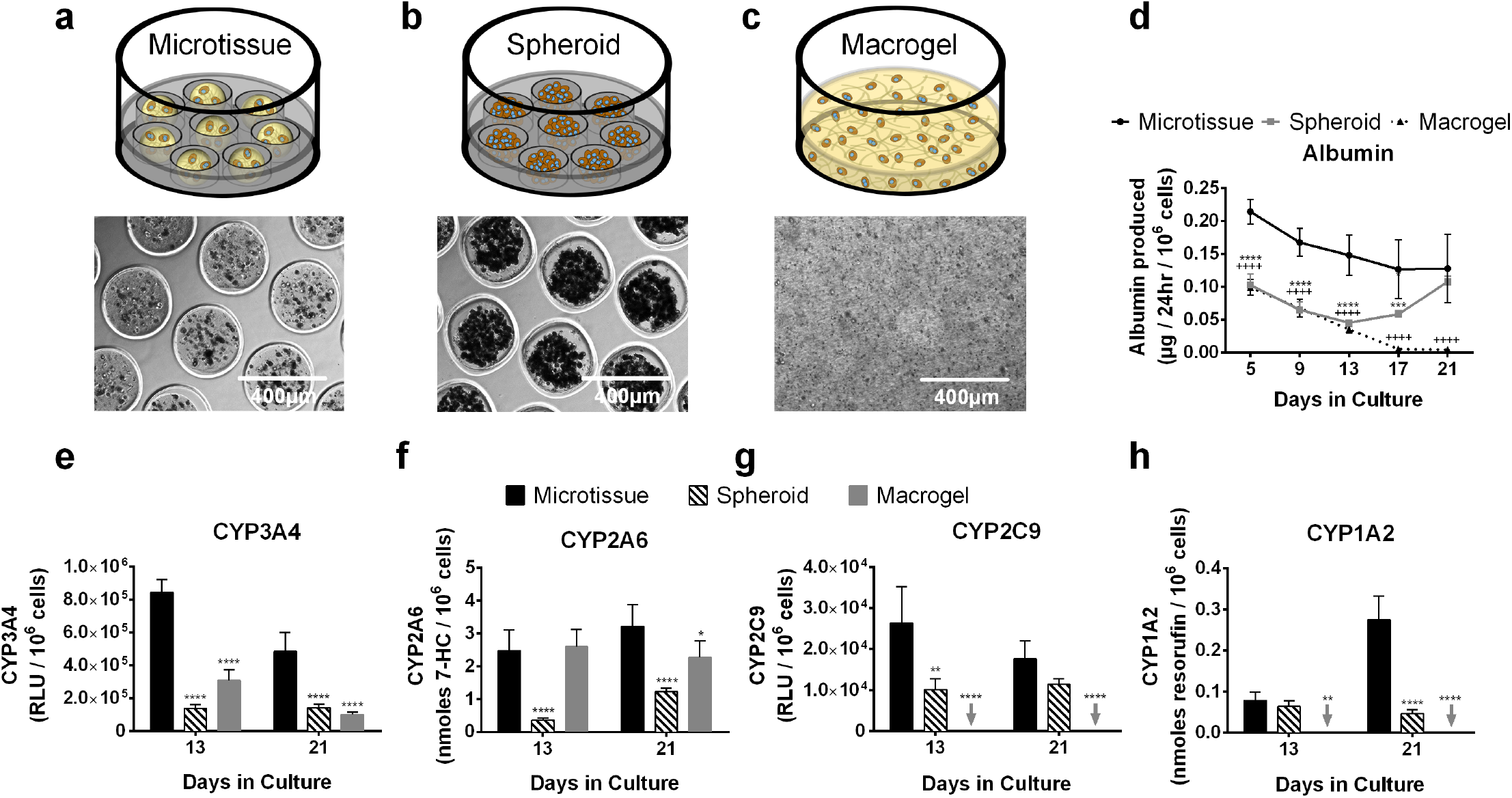
PHH-only microtissues functionally outperform commonly utilized self-assembled spheroids and bulk collagen gels (macrogels). Schematics and phase contrast images for the tested culture models: **(a)** microtissue containing collagen and PHHs formed using the droplet microfluidic device of Fig. 1; **(b)** self-assembled PHH spheroids without any collagen scaffolding; **(c)** macrogels in which the PHH+collagen mixture is dispensed directly into multi-well plates without being subjected to a droplet microfluidic device. **(d)** Albumin production from PHH- only microtissues, spheroids, and macrogels. Statistical significance is displayed for microtissues relative to spheroids (\**p* ≤ 0.05, \*\*\**p* ≤ 0.001, and \*\*\*\**p* ≤ 0.0001) and for microtissues relative to macrogels (++++*p* ≤ 0.0001). Activities of different CYP450 isoenzymes, **(e)** CYP3A4, **(f)** CYP2A6, **(g)** CYP2C9, and **(h)** CYP1A2 in PHH-only microtissues, spheroids and macrogels. Statistical significance is displayed relative to microtissues at the same time point. \**p* ≤ 0.05, \*\**p* ≤ 0.01, and \*\*\*\**p* ≤ 0.0001. Arrows indicate undetectable levels for the indicated function and indicated culture model.

PHHs in microtissues secreted albumin at ~2-fold higher rates than spheroids even with 10-fold fewer cells/volume ratio than the spheroids; similarly, PHHs in microtissues secreted albumin at 4.2-fold and 30-fold higher rates than macrogels after 13 and 21 days in culture, respectively (**Fig. 3d**). Comparably, CYP450 (3A4, 2A6, 2C9, and 1A2) enzyme activities were higher in microtissues than both the spheroids and macrogels. For CYP3A4, microtissues outperformed spheroids by 6.1-fold and 3.4-fold after 13 and 21 days, respectively; similarly, microtissues outperformed macrogels by 2.7-fold and 4.8-fold at the same time-points above (**Fig. 3e**). For CYP2A6, microtissues outperformed spheroids by 6.9-fold and 2.6-fold after 13 and 21 days, respectively; similarly, microtissues outperformed macrogels by 1.4-fold after 21 days (**Fig. 3f**). For CYP2C9, microtissues outperformed spheroids by 2.6-fold and 1.5-fold after 13 and 21 days, respectively; macrogels, on the other hand, had undetectable levels of CYP2C9 activity (**Fig. 3g**). Finally, for CYP1A2, microtissues outperformed spheroids by 1.2-fold and 6-fold after 13 and 21 days, respectively; macrogels, on the other hand, had undetectable levels of CYP1A2 activity (**Fig. 3h**).

Overall, PHH-only spheroids functionally outperformed PHH-only macrogels for CYP1A2/CYP2C9 activities and albumin secretion, while macrogels outperformed spheroids for CYP3A4/CYP2A6 activities. On the other hand, microtissues outperformed both spheroids and macrogels across all measured functions as discussed above.

### 3T3-J2 murine embryonic fibroblasts enhance PHH functions in microtissues at higher levels than primary human HSCs

3T3-J2 fibroblasts are known to enhance PHH functions in both 2D co-cultures ^7^ and 3D self-assembled spheroids ^29^. Therefore, here PHHs and growth-arrested 3T3-J2 fibroblasts (1:1) were either co-encapsulated within the microtissue or fibroblasts were ‘coated’ onto the surface of the collagen-based PHH microtissues and compared to PHH-only microtissues (**Fig. 4a-c**). In contrast to PHH-only microtissues, 3T3-J2 co-encapsulated microtissues and 3T3-J2 coated microtissues were compacted by ~50% due to the fibroblasts. Specifically, 3T3-J2 co-encapsulated microtissues and 3T3-J2 coated microtissues were compacted to an average diameter of 149.2 μm (standard deviation 22.6 μm, n=375 microtissues) and 140.4 μm (standard deviation 13.4 μm, n=375 microtissues), respectively (**Supplemental Fig. 1**).

**Figure 4.**
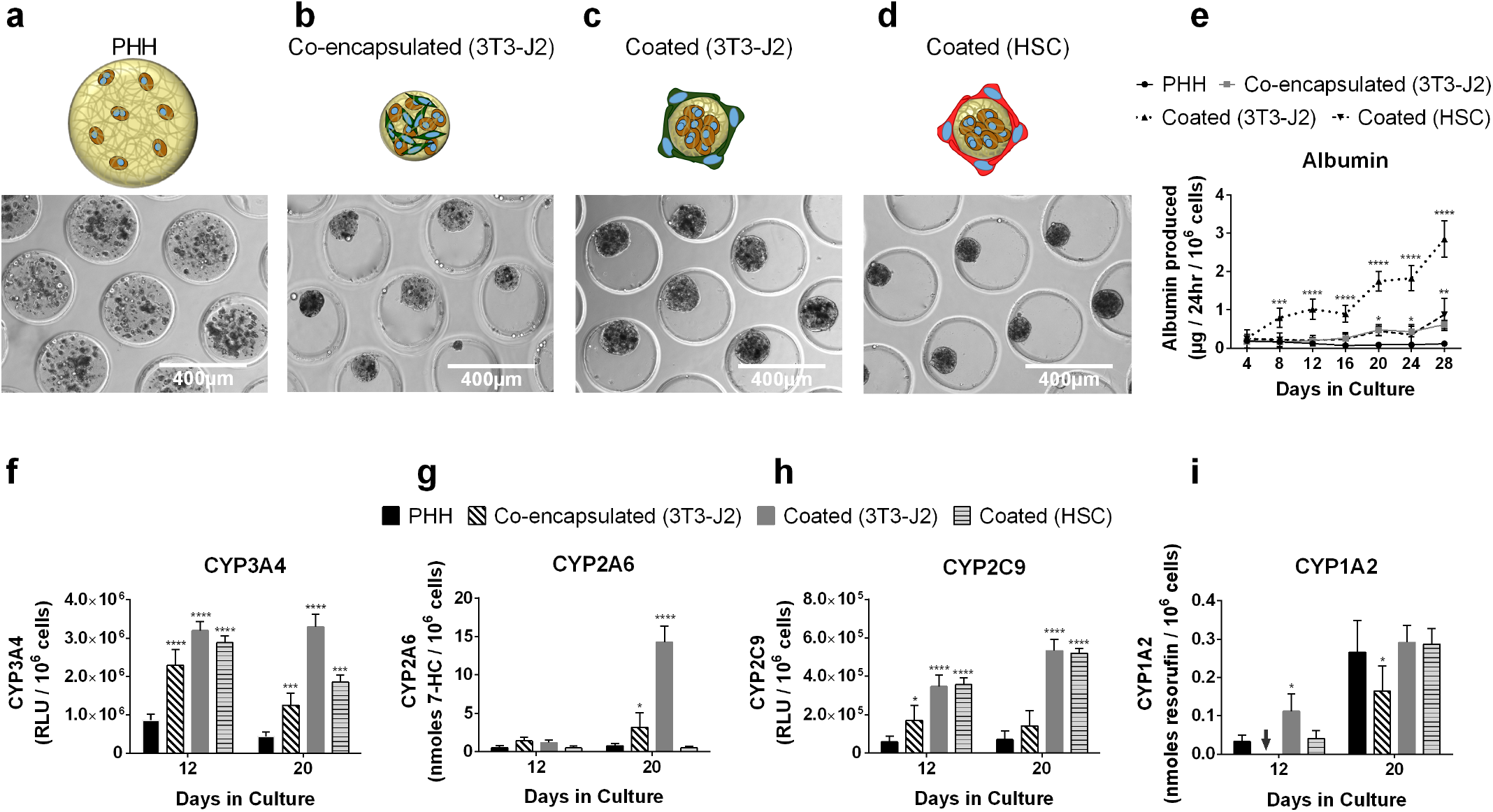
Co-culture of PHHs with either 3T3-J2 mouse embryonic fibroblasts or primary hepatic stellate cells (HSCs) in microtissues enhances liver functions. Phase contrast images for tested culture models: **(a)** Only PHHs encapsulated within collagen microtissues; **(b)** PHHs and 3T3-J2 fibroblasts encapsulated together within the collagen microtissue (‘co-encapsulated (3T3-J2)’); **(c)** 3T3-J2 fibroblasts seeded/coated onto the surface of the polymerized collagen-based PHH microtissues (‘coated (3T3-J2)’); (d) primary hepatic stellate cells (HSCs) seeded/coated onto the surface of the polymerized collagen-based microtissues (‘coated (HSC)’). The smaller size of the microtissues with co-culture indicate compaction of the microtissues by the fibroblasts. **(e)** Albumin secretions from PHH-only, PHH/3T3-J2 co-encapsulated, PHH/3T3-J2 coated, and PHH/HSC coated microtissues. Activities of different CYP450 isoenzymes, **(f)** CYP3A4, **(g)** CYP2A6, **(h)** CYP2C9, **(i)** CYP1A2 in PHH-only, PHH/3T3-J2 co-encapsulated, PHH/3T3-J2 coated, and PHH/HSC coated microtissues. Statistical significance is displayed relative to PHH-only microtissues (\**p* ≤ 0.05, \*\**p* ≤ 0.01, \*\*\**p* ≤ 0.001, \*\*\*\**p* ≤ 0.0001). Arrow indicates undetectable level for the indicated function and indicated culture model.

At a functional level, albumin secretion rates were ~5-fold higher in the 3T3-J2 co-encapsulated microtissues than PHH-only microtissues after 20 days in culture (**Fig. 4e)**. For CYP3A4, 3T3-J2 co-encapsulated microtissues outperformed PHH-only microtissues by 3-fold after 12 and 20 days (**Fig. 4f**). For CYP2A6, 3T3-J2 co-encapsulated microtissues outperformed PHH-only microtissues by 2.8- and 4.4-fold after 12 and 20 days, respectively (**Fig. 4g**). For CYP2C9, 3T3-J2 co-encapsulated microtissues outperformed PHH-only microtissues by 2.7- and 4-fold after 12 and 20 days, respectively (**Fig. 4h**). However, for CYP1A2, PHH-only microtissues outperformed 3T3-J2 co-encapsulated microtissues by 1.6-fold after 20 days (**Fig. 4i**). Fibroblasts were found to be devoid of any of the measured liver functions, including albumin secretion, urea synthesis, and CYP450 activities (data not shown).

3T3-J2 coated microtissues functionally outperformed both 3T3-J2 co-encapsulated and PHH- only microtissues. For albumin, 3T3-J2 coated microtissues outperformed co-encapsulated ones by 5- and 4.2-fold after 8 and 24 days, respectively; similarly, 3T3-J2 coated microtissues outperformed PHH-only microtissues by 4.2- and 20.6-fold at the same time-points above (**Fig. 4e**). For CYP3A4, 3T3-J2 coated microtissues outperformed co-encapsulated ones by 1.3- and 2.7-fold after 12 and 20 days, respectively; similarly, 3T3-J2 coated microtissues outperformed PHH-only microtissues by 3.8- and 8-fold at the same time-points above (**Fig. 4f**). For CYP2A6, 3T3-J2 coated microtissues outperformed co-encapsulated ones by 4.5-fold after 20 days; similarly, 3T3-J2 coated microtissues outperformed PHH-only microtissues by 2.4- and 19.5-fold after 12 and 20 days, respectively (**Fig. 4g**). For CYP2C9, 3T3-J2 coated microtissues outperformed co-encapsulated ones by 2- and 3.7-fold after 12 and 20 days, respectively; similarly, 3T3-J2 coated microtissues outperformed PHH-only microtissues by 6- and 7.6-fold at the same time-points above (**Fig. 4h**). Lastly, for CYP1A2, 3T3-J2 coated microtissues outperformed co-encapsulated ones by 1.8-fold after 20 days; similarly, 3T3-J2 coated microtissues outperformed PHH-only microtissues by 3.4-fold after 12 days though functions across the two models were similar after 20 days (**Fig. 4i**).

Primary HSCs differentiated into myofibroblasts via passaging onto tissue culture plastic have been shown to promote PHH function in 2D culture, albeit at lower levels than 3T3-J2 fibroblasts ^23^. Therefore, here HSCs were coated onto PHH microtissues to determine functional effects in 3D (**Fig. 4d**). HSC-coated microtissues had 5- and 7.4-fold higher albumin secretion rates than PHH-only microtissues after 20 and 28 days in culture, respectively; however, 3T3-J2-coated microtissues had 3.9- and 3.3-fold higher albumin secretion rates than HSC-coated microtissues at the same time-points above (**Fig. 4e**). For CYP3A4, HSC-coated microtissues outperformed PHH-only microtissues by 3.3- and 4.2-fold after 12 and 20 days, respectively; however, 3T3-J2-coated microtissues outperformed HSC-coated microtissues by 1.8-fold after 20 days (**Fig. 4f**). For CYP2A6, unlike 3T3-J2-coated microtissues which upregulated enzymatic activity relative to PHH-only microtissues by up to 19.5-fold on day 20, HSC-coated microtissues did not display enhanced CYP2A6 activity relative to PHH-only microtissues (**Fig. 4g**). For CYP2C9, similar to 3T3-J2-coated microtissues, HSC-coated microtissues outperformed PHH-only microtissues by 6.2- and 7.5-fold after 12 and 20 days, respectively (**Fig. 4h**). Lastly, for CYP1A2, HSC-coated microtissues had similar enzymatic activity as compared to PHH-only microtissues, whereas the 3T3-J2-coated microtissues showed a transient upregulation after 12 days in culture relative to HSC-coated microtissues (**Fig. 4i**).

Overall, 3T3-J2-coated microtissues functionally outperformed HSC-coated microtissues in albumin secretion and three out of the four CYP enzyme activities measured (CYP3A4, CYP2A6, CYP1A2). Therefore, 3T3-J2-coated microtissues were utilized for subsequent studies.

### 3T3-J2-coated microtissues functionally outperform similarly coated self-assembled spheroids and coated macrogels

As with coated microtissues (**Fig. 5a**), 3T3-J2 fibroblasts were coated onto self-assembled PHH spheroids (1:1 ratio between the two cell types) (**Fig. 5b**) and such coated spheroids were also compacted by ~35% due to the fibroblasts with an average diameter of 133.7 μm (standard deviation 21.6 μm, n=375 microtissues); however, coated microtissues had a more consistent size distribution than coated spheroids (**Supplemental Fig. 1**). Similarly, 3T3-J2 fibroblasts were coated onto the surface of PHH macrogels at a 1:1 ratio (**Fig. 5c**).

**Figure 5.**
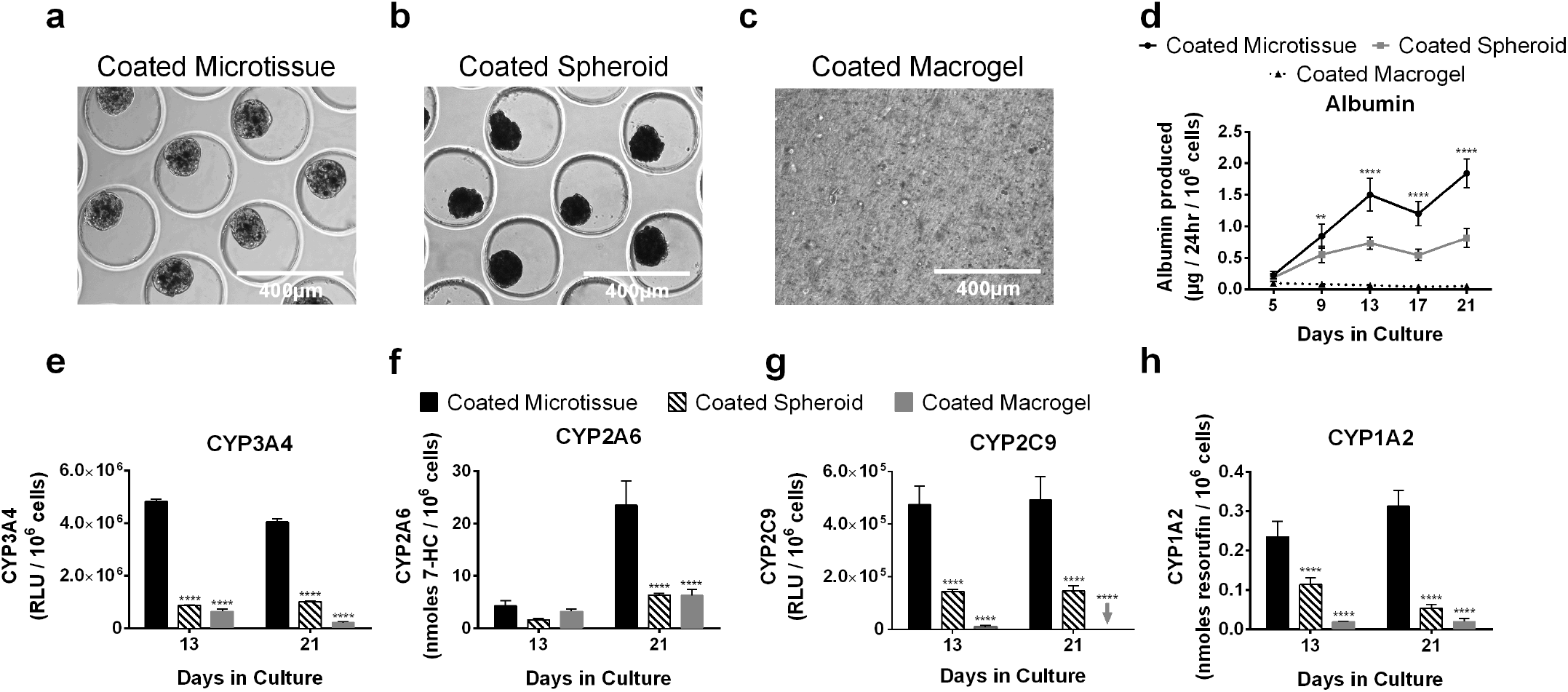
Coated (co-culture) microtissues functionally outperform coated self-assembled spheroids and macrogels. **(a)** 3T3-J2 fibroblasts were seeded/coated onto the surface of the polymerized collagen-based PHH microtissues (‘coated microtissue’). **(b)** Similarly, self-assembled PHH spheroids were coated with 3T3-J2 fibroblasts (‘coated spheroids’) and **(c)** macrogels were coated with 3T3-J2 fibroblasts (‘coated macrogels’). **(d)** Albumin production in the coated microtissues, coated spheroids, and coated macrogels. Statistical significance is displayed for coated microtissues relative to coated spheroids at the same time point (\*\**p* ≤ 0.01, and \*\*\*\**p* ≤ 0.0001). Activities of different CYP450 isoenzymes, **(e)** CYP3A4, **(f)** CYP2A6, **(g)** CYP2C9, and **(h)** CYP1A2 in the coated microtissues, coated spheroids, and coated macrogels. Statistical significance is displayed for coated spheroids and coated macrogels relative to coated microtissues at the same time point (\*\*\*\**p* ≤ 0.0001). Arrow indicates undetectable level for the indicated function and indicated culture model.

At a functional level, coated microtissues outperformed both coated spheroids and coated macrogels. For albumin secretion, coated microtissues outperformed coated spheroids by 2-fold and 2.3-fold after 13 and 21 days, respectively; similarly, coated microtissues outperformed coated macrogels by 22.9-fold and 36.2-fold at the same time-points above (**Fig. 5d**). For CYP3A4, coated microtissues outperformed coated spheroids by 5.6-fold and 4-fold after 13 and 21 days, respectively; similarly, coated microtissues outperformed coated macrogels by 7.6-fold and 19.1-fold (**Fig. 5e**). For CYP2A6, coated microtissues outperformed coated spheroids by 2.5-fold and 3.7-fold after 13 and 21 days, respectively; similarly, coated microtissues outperformed coated macrogels by 1.3-fold and 3.7-fold (**Fig. 5f**). For CYP2C9, coated microtissues outperformed coated spheroids by 3.3-fold and 3.4-fold after 13 and 21 days, respectively; similarly, coated microtissues outperformed coated macrogels by 54-fold after 13 days, while CYP2C9 activity was undetectable in macrogels after 21 days (**Fig. 5g**). Finally, for CYP1A2, coated microtissues outperformed coated spheroids by 2-fold and 5.8-fold after 13 and 21 days, respectively; similarly, coated microtissues outperformed coated macrogels by 12-fold and 15.7-fold (**Fig. 5h**).

Overall, coated spheroids functionally outperformed coated macrogels, with CYP2A6 being the sole exception. On the other hand, coated microtissues outperformed both coated spheroids and coated macrogels.

### 3T3-J2-coated microtissues display high levels of liver functions and gene expression for 6 weeks

Coated microtissues were assessed for functions over 6 weeks and these functions were compared to functions in PHHs within 2-4 hours of thawing (day 0) as the gold standard for short-term (acute) assays before PHHs severely decline in functions in conventional 2D mono-culture formats ^5, 13^. Albumin secretion was minimal in PHHs on day 0 but ramped up over the first 3 weeks in coated microtissues to ~6μg/day/million PHHs and remained relatively stable until day 41 when the cultures were sacrificed (**Fig. 6a**). Urea secretion was undetectable in PHHs on day 0 but increased in coated microtissues over the first 9 days to ~57-g/day/million PHHs and remained statistically stable for 41 days (**Fig. 6b**). CYP3A4 activity in the coated microtissues was ~9% of day 0 activity after 9 days and increased to ~20% to 32% for the remaining culture duration (**Fig. 6c**). CYP2A6 activity in the coated microtissues was ~15% of day 0 activity after 9 days and increased to ~113% to 183% for the remaining duration (**Fig. 6d**). CYP2C9 activity in the coated microtissues was ~36% of day 0 activity after 9 days and decreased to ~21% to 26% for the remaining culture duration (**Fig. 6e**). Lastly, CYP1A2 activity in the coated microtissues was ~70% of day 0 activity after 9 days and increased to ~100% to 153% for the remaining duration (**Fig. 6f**).

**Figure 6.**
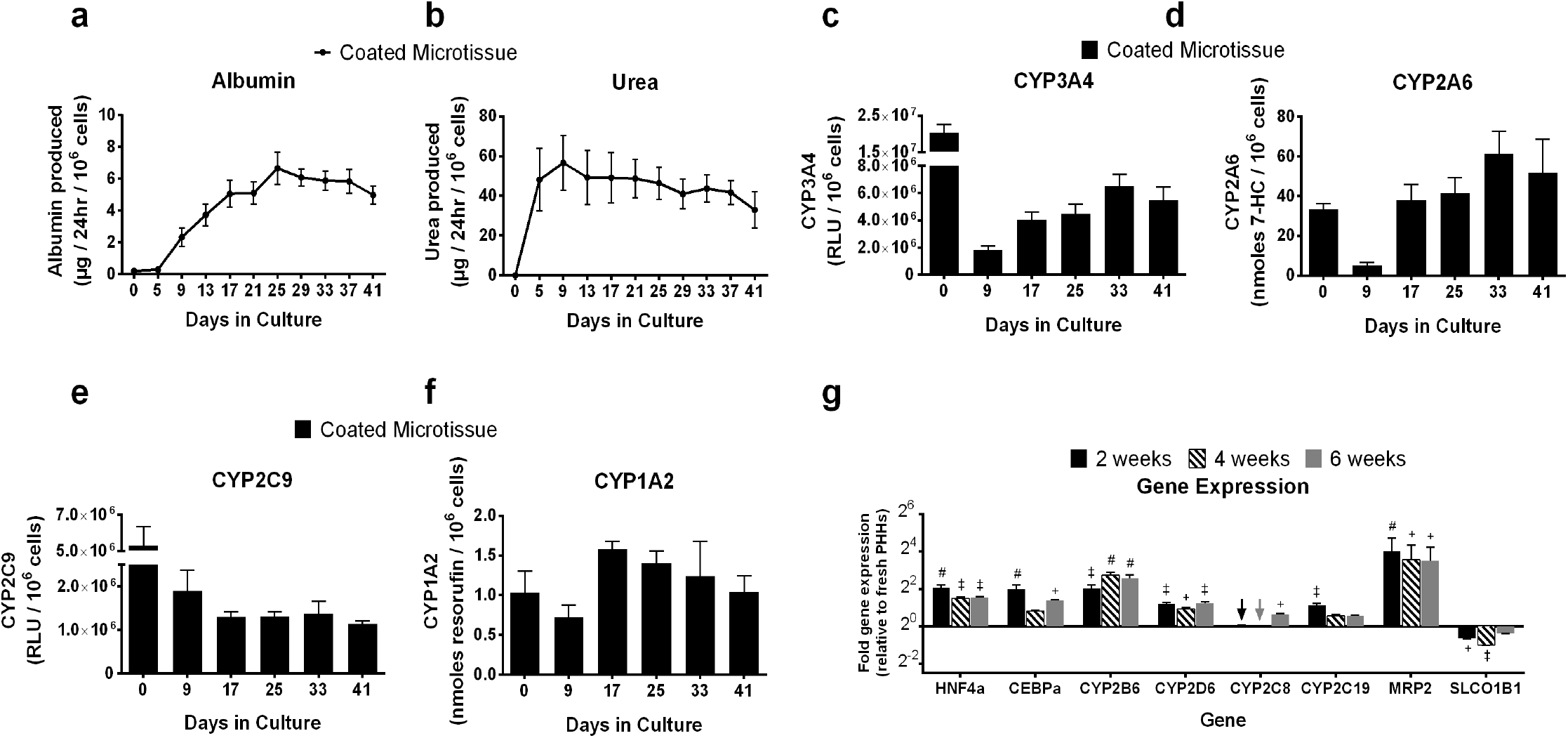
Coated (co-culture) microtissues display stable levels of liver functions and gene expression for 6 weeks. 3T3-J2 fibroblasts were seeded/coated onto the surface of the polymerized collagen-based PHH microtissues (‘coated microtissue’) and cultured for ~6 weeks. **(a)** Albumin production, **(b)** urea production, and the activities of different CYP450 isoenzymes, **(c)** CYP3A4 activity, **(d)** CYP2A6, **(e)** CYP2C9, and **(f)** CYP1A2 in coated microtissues over time. **(g)** Quantitative gene expression of hepatic transcription factors (*HNF4a, CEBPa)*, CYP450 enzymes (*CYP2B6, CYP2D6, CYP2C8, CYP2C19)*, canalicular transporter (*MRP2),* and basolateral transporter (*SLCO1B1)* in coated microtissues at 2, 4, and 6 weeks as compared to freshly thawed PHHs (line 2^0^) that were immediately lysed in suspension for RNA (i.e. expression levels in the coated microtissues at the 2^0^ line are near identical to the levels in freshly thawed PHHs). Arrows indicate detectable levels at the 2^0^ line. Statistical significance for gene expression is displayed for coated microtissues at 2, 4, and 6 weeks relative to freshly thawed PHHs (+*p* ≤ 0.01, ‡*p* ≤ 0.001, #*p* ≤ 0.0001).

Gene expression levels of major liver-specific markers, including master transcription factors (*HNF4a* and *CEBPa*), CYP450 enzymes (*CYP2B6, CYP2D6, CYP2C8,* and *CYP2C19*), and drug transporters (*MRP2* and *SLCO1B1*) were characterized in the coated microtissues after 2, 4, and 6 weeks following culture initiation (**Fig. 6g**). Gene expression in the coated microtissues was normalized to human *GAPDH* and further normalized to gene expression data in PHHs immediately following thawing (day 0). The gene expression levels of 7 of the 8 genes were within 2-fold or higher in the coated microtissues than the levels in freshly thawed PHHs (e.g. after 6 weeks, 2.9-, 2.6-, 6-, 2.4-, 1.6-, 1.5-, and 11.4-fold higher levels in microtissues than freshly thawed PHHs for *HNF4a*, *CEBPa*, *CYP2B6, CYP2D6, CYP2C8, CYP2C19,* and *MRP2,* respectively). In contrast, *SLCO1B1* level was downregulated in the coated co-culture microtissues than the freshly thawed PHHs (e.g. after 6 weeks, 77% retention in microtissues of freshly thawed PHH levels).

### 3T3-J2-coated microtissues respond in a clinically-relevant manner to drug-mediated CYP induction and hepatotoxicity

Coated microtissues were allowed 9 days to reach steady state levels of functions and then treated with rifampin or omeprazole for a total of 4 days with fresh drug added at the 2-day medium exchange. CYP3A4 and CYP2C9 activities were assessed in the rifampin-treated cultures whereas CYP1A2 activity was assessed in the omeprazole-treated cultures; DMSO-treated control cultures were used calculate fold changes in induction of CYP activities due to drug treatment. Rifampin caused ~7-fold induction in CYP3A4 activity and ~5-fold induction in CYP2C9 activity relative to the DMSO control, while omeprazole caused ~2-fold induction in CYP1A2 activity (**Fig. 7a**).

**Figure 7.**
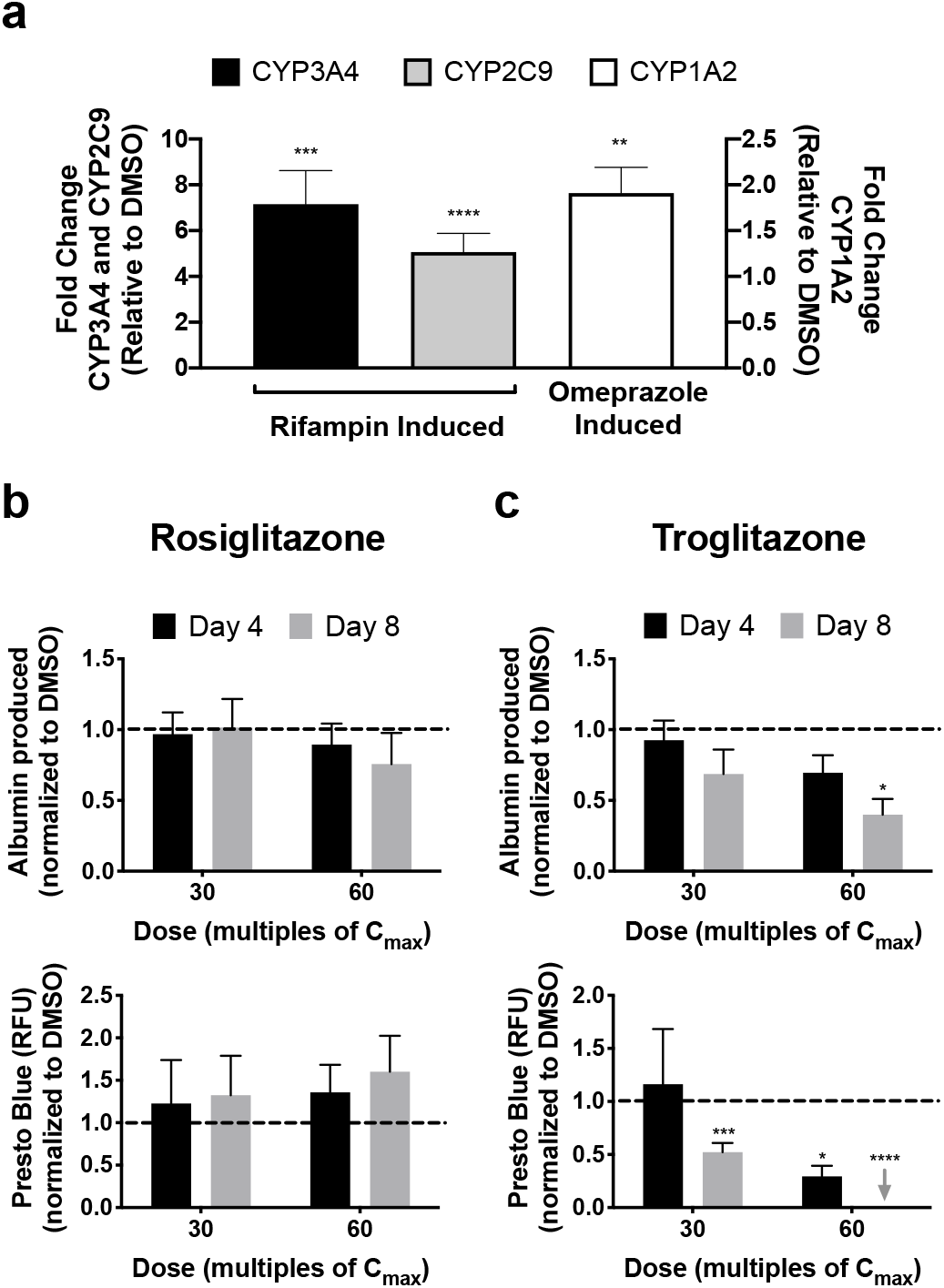
Coated (co-culture) microtissues can be used for drug-mediated CYP induction and hepatotoxicity assessments. 3T3-J2 fibroblasts were seeded/coated onto the surface of the polymerized collagen-based PHH microtissues (‘coated microtissues’). Coated microtissues were allowed ~9 days to functionally stabilize and then treated with rifampin or omeprazole for a total of 4 days with fresh drug added to the culture medium at the 2-day medium exchange. **(a)** CYP3A4 and CYP2C9 activities in coated microtissues treated with rifampin, and CYP1A2 activity in the coated microtissues treated with omperazole. Enzyme activities in the drug-treated microtissues were normalized to enzyme activity in microtissues treated with DMSO alone. Statistical significance is displayed relative to DMSO-treated controls (\*\**p* ≤ 0.01, \*\*\**p* ≤ 0.001, and \*\*\*\**p* ≤ 0.0001). Additionally, coated microtissues were allowed ~9 days to functionally stabilize and then treated with rosiglitazone or troglitazone at two concentrations for each drug, 30× and 60× C_max_ (C_max_ = maximum drug concentration measured in human plasma; C_max_ for troglitazone= 2.82 μg/mL and C_max_ for rosiglitazone = 0.373 μg/mL), for a total of 8 days with fresh drug added to culture medium at the 4-day medium exchange. **(b)** Albumin secretion, specific to PHHs (top graph), and overall co-culture viability (bottom graph, PrestoBlue^TM^ conversion to the metabolite, resazurin) in coated microtissues treated with rosiglitazone. Data in drug-treated coated microtissues were normalized to the corresponding data in DMSO-only treated cultures. **(c)** Similar graphs as panel (a) except coated microtissues were treated with troglitazone. For drug toxicity data, statistical significance is relative to DMSO-treated control cultures. \**p* ≤ 0.05, \*\*\**p* ≤ 0.001, and \*\*\*\**p* ≤ 0.0001.

Coated microtissues stabilized over 9 days were treated with two drugs for type 2 diabetes mellitus, rosiglitazone or troglitazone, at two concentrations for each drug, 30× and 60× C_max_ (C_max_: maximum drug concentration measured in human plasma), for a total of 8 days with fresh drug added at the 4-day medium exchange. Rosiglitazone is a non-hepatotoxin, while troglitazone was withdrawn from the market due to severe hepatotoxicity ^30^. Albumin secretion (hepatocyte marker) and overall co-culture viability (metabolism of PrestoBlue™) were assessed after 4 and 8 days of drug treatment, and the data were normalized to the data in DMSO-only treated cultures. No statistically significant downregulation of either albumin or viability was observed in coated microtissues treated with rosiglitazone (**Fig. 7b**). In contrast, coated microtissues treated with troglitazone displayed time- and concentration-dependent decreases in albumin and viability (**Fig. 7c**). Albumin secretion declined to 92% and 69% of DMSO-only controls after treatment of coated microtissues with 30× C_max_ of troglitazone for 4 and 8 days, respectively, whereas it declined to 69% and 40% after treatment with 60× C_max_ at the same time-points above. Similarly, viability declined to 29% of DMSO controls after treatment of coated microtissues with 30× C_max_ of troglitazone for 8 days, whereas it declined to 52% after treatment with 60× C_max_ for 4 days and was undetectable following treatment with this concentration for 8 days.

## DISCUSSION

In contrast to a limited application scope of 2D models, human liver models that are 3D in architecture have utility not only for preclinical drug development but also for elucidating the reorganization of cell-cell and cell-ECM interactions in liver disease and as building blocks for cell-based therapies in the clinic. In the absence of a functional vasculature, it is important to restrict the size of the 3D human liver models to ~300 μm in diameter to allow for adequate oxygen and nutrient transfer. While such miniaturized models can be created using specialized plates that allow spheroid generation, typically the throughput of such approaches is limited. In contrast, droplet microfluidics provides the ability to rapidly create highly reproducible microtissues containing one or more cell types; however, the use of this technology has been restricted to rat hepatocytes or cancerous hepatic cell lines so far. Thus, here we utilized droplet microfluidics to create 3D human liver microtissues within a natural collagen-based hydrogel microenvironment that displayed higher PHH functions than commonly employed self-assembled spheroids and bulk hydrogels (macrogels); hepatic functions were enhanced and stabilized for up to 6 weeks in the presence of supportive fibroblasts, though 3T3-J2 fibroblasts induced higher functions in PHHs than primary human HSCs. Placement of the microtissues into agarose microwells cast within multi-well plates prevented the aggregation of the microtissues and allowed probing with prototypical drugs for CYP induction and hepatotoxicity assays.

Human liver microtissues were compared to self-assembled spheroids and macrogels that are routinely used to access 3D liver biology for structure-function studies and drug testing. All culture models enabled PHH functions for 2 weeks, confirming that a 3D microenvironment can sustain PHH functions over rapidly declining 2D monocultures on collagen-I adsorbed onto plastic dishes ^5, 13^. However, the microtissues displayed significantly higher albumin and CYP450 enzyme activities than spheroids and macrogels. CYP450 enzymes metabolize >50% of marketed drugs and their expression is influenced by age, sex, genetic polymorphisms, drugs/chemicals, hormones/cytokines, and disease states ^31^. Thus, it is critical to maintain physiologically-relevant levels of these enzymes *in vitro* towards elucidating *in vivo*-like compound metabolism and toxicity ^5, 13^; our results suggest that microtissues may enable such an outcome over the gold standard spheroids and macrogels.

The higher CYP450 activities in microtissues than spheroids and macrogels may be due to a more optimal and consistent microenvironment around PHHs. First, microtissues allow for the ligation of PHH integrins to the collagen immediately as opposed to the time-dependent ligation with cell-secreted ECM in the spheroids. Second, microtissues maintain a consistent inter- and intra-donor ECM microenvironment as opposed to the likely variable ECM secretion rates by different donors in spheroids. Third, the microscale dimensions of microtissues likely facilitate the optimal delivery of oxygen and nutrients than the macrogels. Lastly, homotypic cell-cell interactions, which have been shown to be critical for PHH functions and polarity ^5^, are likely better facilitated within the microtissues than the macrogels at similar cell seeding densities. Therefore, microtissues may provide a more reproducible platform for drug development than spheroids.

In addition to generating higher liver-specific function, microtissues also improve the logistical and cost considerations for screening compounds over spheroids and macrogels. While microtissues were found to form robustly here with multiple PHH donors, spheroids are difficult to stably form with >50% of PHH donors ^10^, which may be due to differential ECM secretion rates across PHH donors. Furthermore, microtissues use ~10% of PHHs used in spheroids, but still enable the highest CYP450 activities when normalized to cell numbers. Lastly, the fabrication of macrogels requires large amounts of expensive ECM and thus places considerable limits on the sample size for each condition; in contrast, hundreds of microtissues can be tracked for each condition. Therefore, PHH microtissues can allow the screening of large compound libraries or various disease stimuli with multiple PHH lots towards identifying lead molecules/pathways for further testing, either *in vitro* or *in vivo*.

Co-culture with liver- and non-liver-derived non-parenchymal cell (NPC) types can affect hepatocyte functions in the developing and adult livers ^5, 13^. Some of these interactions can be replicated *in vitro* by co-culturing primary hepatocytes with NPCs, such as rat hepatocytes co-cultured with either C3H/10T1/2 mouse embryo cells ^32^, 3T3-J2 murine embryonic fibroblasts ^33^, or human fibroblasts ^34^. Of these, 3T3-J2 fibroblasts express molecules present in the liver (e.g. decorin, VEGF, and T-cadherin) at higher levels than other 3T3 fibroblast clones ^35^, which allows 3T3-J2 fibroblasts to induce higher PHH functions in 2D co-cultures ^7^; such an effect requires contact between the cell types as fibroblast-conditioned medium alone does not suffice for stabilizing hepatic functions ^36^. Furthermore, 3T3-J2 fibroblasts induce higher levels of functions in PHHs than human liver sinusoidal endothelial cells ^22^, hepatic stellate cells ^23^, and Kupffer cells ^37^, albeit in 2D culture formats.

Owing to the positive effects of 3T3-J2 fibroblasts on PHH functions, these fibroblasts were co-cultured with PHHs in the microtissues via co-encapsulation of both cell types or by coating the PHH-encapsulated microtissues with fibroblasts; similarly, self-assembled PHH spheroids and PHH-containing collagen macrogels were also coated with fibroblasts to elucidate the role of microscale collagen presentation on PHH functions in the presence of fibroblasts. The fibroblasts enhanced PHH functions over PHH mono-cultures placed within 3D microtissues, spheroids, and macrogels. However, the fibroblast coated microtissues displayed significantly higher functions than both the co-encapsulated microtissues, coated spheroids, and coated macrogels. The higher functions of the coated versus co-encapsulated microtissues may be due to increased PHH homotypic contacts during compaction, which is crucial for PHH polarity and function, whereas co-encapsulated microtissues only enable a randomly dispersed co-culture that has been shown to be insufficient for promoting adequate PHH contacts even in 2D monolayers^7^. Additionally, coated microtissues were more homogenous in size after compaction as compared to co-encapsulated microtissues, which may further enable consistent PHH homotypic contacts and thus higher function. On the other hand, while considerable compaction was also observed in the coated spheroids, the collagen scaffolding within the microtissues was necessary to further enhance PHH functions, suggesting that the increase in ECM density in the coated microtissues may also play a role in increased function.

Primary HSCs, especially those that have been differentiated into myofibroblasts, have been previously shown to support some hepatic functions *in vitro* ^23, 38^ even though they are implicated in causing liver fibrosis due to excessive collagen deposition *in vivo ^39^*. Here, we compared the effects of 3T3-J2 fibroblasts and primary HSCs in induction of PHH functions within the 3D microtissues. HSC-coated microtissues displayed higher albumin secretion, CYP3A4 activity, and CYP2C9 activity than PHH-only microtissues, whereas CYP2A6 and CYP1A2 activities were similar across the two models. However, 3T3-J2-coated microtissues functionally outperformed HSC-coated microtissues for albumin secretion and three out of the four CYP enzyme activities measured (CYP3A4, CYP2A6, CYP1A2), while CYP2C9 activity was similar in both models. These results suggest that the 3T3-J2-coated microtissues may be most suitable for routine assessment of drug responses in highly functional PHHs. Nonetheless, the ability of the HSC-coated microtissues to support PHH phenotype for several weeks *in vitro* suggests that the microtissue platform may be suitable to model PHH-HSC interactions in physiology and disease (e.g. excessive HSC proliferation and ECM deposition/reorganization induced via drugs and/or excess nutritional stimuli as *in vivo*).

In the 3T3-J2-coated microtissues, PHHs displayed long-term and stable phenotypic functions and gene expression for 6+ weeks at comparable levels to freshly thawed PHHs with some noted exceptions. CYP2A6 and CYP1A2 activities, as well as the expression of 8 key liver transcripts, in the coated microtissues recovered to within 2-fold of the levels in freshly thawed PHHs. On the other hand, albumin and urea secretions were not detected in freshly thawed PHHs but reached stable steady-state levels in coated microtissues, which may be due to the rapid degradation of secretory capacities following isolation of PHHs from the liver; indeed, recovery of albumin and urea secretions has also been observed previously in PHH/NPC co-cultures ^7, 29, 40^. Additionally, CYP3A4 and CYP2C9 activities in the coated microtissues were ~25-30% of the activities measured in freshly thawed PHHs. Since CYP3A4 and 2C9 enzyme activities can be induced by several drugs ^41, 42^, the initial presence of any inducer drugs in the donor’s liver and thus PHHs may be a confounding factor in our findings. Nonetheless, the enhanced longevity of coated microtissues can potentially enable the elucidation of the chronic effects of drugs, industrial chemicals, and disease stimuli (e.g. hepatitis B virus, alcohol, and excess dietary triggers) on PHH functions.

The 3T3-J2-coated microtissues were found to display clinically-relevant drug responses. Specifically, omperazaole induced CYP1A2 activity in microtissues, likely due to binding to aryl hydrocarbon receptor ^43^; rifampin induced both CYP3A4 and CYP2C9, likely due to binding to pregnane X receptor ^44^. Fold changes of induction in microtissues were similar to that observed previously in PHH cultures when accounting for donor-to-donor differences ^7^, suggesting that the microtissue platform is suitable for drug-mediated CYP induction assays. In addition to CYP induction, coated microtissues were treated with troglitazone (hepatotoxin) or rosiglitazone (non-hepatotoxin) for 8 days and at concentrations up to 60× C_max_ for each drug since these parameters were previously shown to allow appropriate appraisal of safety margins for diverse pharmaceuticals without causing an increase in the detection of false positives (i.e. loss of specificity) ^21, 25^. Rosiglitazone did not affect either PHH albumin or viability in the coated microtissues, which is consistent with its relatively safe profile with respect to liver liabilities in humans ^30^. In contrast, troglitazone caused a time- and concentration-dependent loss of albumin secretion and cell viability. Co-culture viability was affected more severely than albumin in the supernatants, which may be due to the detection of albumin from lysed PHHs; however, fibroblast contribution to the co-culture viability trends requires further investigation using cell-specific staining markers. Furthermore, an appraisal of the overall sensitivity and specificity of microtissues for preclinical assessment of drug-induced hepatotoxicity would necessitate testing of a larger panel of drugs (>50) from diverse classes and mechanisms of action, which we plan to pursue in follow-up studies. Nonetheless, the proof-of-concept data here suggests that 3T3-J2 fibroblast coating does not inhibit the utility of microtissues for investigating PHH responses to compounds following repeat exposure.

Our modular microtissues can be further tuned to investigate other features of the PHH microenvironment in the liver. First, collagen I can be augmented with additional liver-inspired ECM molecules such as other collagens (e.g. collagen IV), fibronectin, laminin, and proteoglycans (e.g. decorin). Second, custom microtissues containing liver NPCs, such as liver sinusoidal endothelial cells, hepatic stellate cells, and Kupffer cells/macrophages, can be created to test specific hypotheses around PHH-NPC interactions in physiology and disease. Lastly, microfluidic devices can be used to induce gradients of oxygen and hormones across microtissues to mimic differential PHH functions as in liver zonation ^45^, which can be useful to determine whether identified toxic compounds show a zonal preference ^46^.

In conclusion, we showed that 3D collagen-I microtissues containing PHHs display phenotypic stability, including CYP450 enzyme activities, for at least 6 weeks within multi-well plates without the need for any fluid perfusion. PHH functions in such microtissues can be further enhanced via coated co-culture with primary human HSCs or 3T3-J2 fibroblasts, with the latter providing for the highest levels for the majority of PHH functions. Finally, we showed that 3T3-J2-coated 3D microtissues are suitable to assess drug-mediated CYP induction and hepatotoxicity following repeat drug exposure. Ultimately, the functionally optimized microtissues could serve to reduce drug attrition, enable the screening of molecules to optimize cell functions in cell-based therapies, and aid in the studies of liver physiology and disease.

## MATERIALS AND METHODS

### Microfluidic and microwell device fabrication

Polydimethylsiloxane (PDMS)-based microfluidic devices consisting of a single emulsion droplet generator with 300 μm straight channel and 150 μm nozzle were fabricated, bonded to glass slides, and coated with hydrophobic Novec™ 1720 (3M, Maplewood, MN) using published procedures ^15^. Molten agarose (2% w/v) was prepared and dispensed into each well of a 24-well tissue culture polystyrene plate. PDMS stamps with ~1000 micropillars were placed into the molten agarose within each well of a 24-well polystyrene plate. After the agarose cooled, the PDMS stencils were removed, forming ~300 μm diameter × ~300 μm deep wells (microwells) in the casted agarose ^16^. PDMS devices and agarose microwells were sterilized via autoclaving or 70% ethanol treatment (1 hour), respectively.

### Primary human hepatocyte (PHH) mono-cultures

A solution of rat tail collagen, type I (Corning Life Sciences, Tewksbury, MA) in acetic acid was first diluted in 1X phosphate buffered saline (PBS, Corning) to 6 mg/mL on ice, and then the pH was neutralized to 7.4-7.6 using 1N NaOH. Cryopreserved PHH were thawed, counted, and viability (>85%) was assessed as previously described ^21^. PHHs were resuspended in the neutral collagen solution at varying densities (1.25- to 5e6 cells/mL). The collagen solution containing cells was perfused into the microfluidic device inlet at 150 μL/hour in a cold room (4ºC) while fluorocarbon oil (FC-40, Sigma-Aldrich) with 2% 008-fluoroSurfactant (RAN Biotechnologies, Beverly, MA) was perfused at 650 μL/hour at the T-junction to produce PHH/collagen emulsions (herein referred to as ‘microtissues’). Microtissues were collected in a 1.5 mL tube that was heated at 37°C to promote collagen polymerization. Polymerized microtissues were rinsed, resuspended in culture medium, counted, and seeded into the agarose microwells within a 24-well plate (~600 microtissues/well). Hepatocyte culture medium, the composition of which was described previously ^22^, was replaced on microtissues every 4 days (400 μL/well). Lastly, conventional PHH mono-cultures were created by either seeding PHHs directly into agarose microwells (200K cells in 400 μL/well in a 24-well plate) to form self-assembled spheroids, or by seeding a PHH and collagen (6 mg/mL) mixture (20K cells in 200 μL/well of collagen) directly in a 24-well plate to form bulk gels (herein referred to as ‘macrogels’).

### Co-cultures of PHHs with 3T3-J2 murine embryonic fibroblasts or primary hepatic stellate cells (HSC)

3T3-J2 fibroblasts and primary HSCs were passaged onto tissue culture plastic as previously described ^23^; the HSCs become myofibroblasts when passaged *in vitro*. Both fibroblast cell types were subsequently growth arrested by incubating with 1 μg/mL mitomycin-C (Sigma-Aldrich, St. Louis, MO) in culture medium for 4 hours prior to detachment from the culture substrates using trypsin as previously described ^7^. Microtissues were created as described above except that the PHHs and 3T3-J2 fibroblasts at a 1:1 ratio were first co-suspended in the collagen solution prior to injection into the microfluidic device to create ‘co-encapsulated’ microtissues. Alternatively, PHH-only microtissues were created, placed into agarose microwells within a 24-well plate, and then 3T3-J2 fibroblasts or HSCs were seeded onto the polymerized PHH microtissues at ~1:1 PHH:fibroblast or HSC ratio; the fibroblasts or HSCs preferentially attached to the collagen microtissues as opposed to the non-adhesive agarose to create so-called ‘coated’ co-culture microtissues. Culture medium was replaced as described above for PHH mono-cultures.

Additionally, self-assembled PHH / 3T3-J2 spheroids were created by seeding PHHs into agarose microwells as described above and then seeding 3T3-J2 fibroblasts the next day at a 1:1 ratio with the PHHs (200K cells/well for each cell type in a 24-well plate). Finally, PHH-containing macrogels were created as described above and then 3T3-J2 fibroblasts were seeded onto the surface of the polymerized PHH macrogels the next day at a 1:1 ratio with the PHHs (20K cells/well for each cell type in a 24-well plate).

### Hepatocyte functional assessments

Culture supernatants were assayed for albumin using a sandwich enzyme-linked immunosorbent assay (ELISA, Bethyl Laboratories, Montgomery, TX) with horseradish peroxidase detection and 3,3’,5,5’-tetramethylbenzidine (TMB, Rockland Immunochemicals, Boyertown, PA) as the substrate ^7^. Urea concentration in supernatants was assayed using a colorimetric end-point assay utilizing diacetyl monoxime with acid and heat (Stanbio Labs, Boerne, TX) ^7^. Absorbance values were quantified on the Synergy H1 multi-mode plate reader (BioTek, Winooski, VT).

CYP3A4 and CYP2C9 enzyme activities were measured by incubating the cultures with luciferin-IPA or luciferin-H substrates (Promega Life Sciences, Madison, WI) for 3 hours, respectively. The metabolite, luciferin, was quantified via luminescence detection on the Synergy H1 multi-mode plate reader according to manufacturer’s protocols. CYP1A2 and CYP2A6 activities were measured by incubating the cultures with 5 μM 7-ethoxyresorufin or 50 μM coumarin (Sigma-Aldrich) for 3 hours, respectively. The metabolites, resorufin and 7-hydroxycoumarin (7-HC), generated from 7-ethoxyresorufin and coumarin, respectively, were quantified via fluorescence detection (excitation/emission: 550/585 nm for resorufin, and 355/460 nm for 7-HC) on the Synergy H1 multi-mode plate reader ^24^.

### Gene expression analysis

Total cellular RNA was extracted using TRIzol™ (Thermo Fisher Scientific, Waltham, MA), purified using RNeasy mini kit (Qiagen, Germantown, MD), and genomic DNA was digested using Optizyme™ recombinant DNase-I digestion kit (Thermo Fisher) per manufacturers’ instructions. Purified RNA was then reverse transcribed into complementary DNA (cDNA) using the high-capacity cDNA reverse transcription kit (Thermo Fisher) per manufacturer’s instructions on a MasterCycler RealPlex 2 (Eppendorf, Hauppauge, NY). Then, 250 ng of cDNA was added to each quantitative polymerase chain reaction (qPCR) along with the Taqman™ master mix (Thermo Fisher) and pre-designed Taqman human-specific primer/probe sets per manufacturer’s protocols. The primer/probe sets were selected to be human-specific without cross-reactivity to mouse DNA, and included glyceraldehyde 3-phosphate dehydrogenase (*GAPDH*), hepatocyte nuclear factor 4-alpha (*HNF4a)*, CCAAT/enhancer binding protein alpha (*CEBPa*), cytochrome P450 2B6 (*CYP2B6)*, cytochrome P450 2D6 (*CYP2D6)*, cytochrome P450 2C8 (*CYP2C8*), cytochrome P450 2C19 (*CYP2C19*), multidrug resistance associated protein 2 (*MRP2*), and solute carrier organic anion transporter family member 1B1 (*SLCO1B1*). Taqman primer/probe sequences are proprietary to the manufacturer. Hepatic gene expression was normalized to *GAPDH*.

### Drug-mediated CYP induction and drug-induced hepatotoxicity studies

PHH/3T3-J2 coated microtissues were allowed 9 days to functionally stabilize and then treated with either 6.25 μM rifampin or 3.125 μM omperazole (Sigma-Aldrich) for a total of 4 days with fresh drug added at the 2-day medium exchange. The drug was dissolved in 100% dimethylsulfoxide (DMSO, Corning Life Sciences). The final DMSO concentration in the culture medium was kept at 0.1% (v/v), and a DMSO-only control culture was used to calculate fold changes due to the drug treatment. After 4 days of drug treatment, CYP3A4 and CYP2C9 activities were measured for rifampin-treated cultures and CYP1A2 activity was measured for omeprazole-treated cultures as described above.

PHH/3T3-J2 coated microtissues were allowed 9 days to functionally stabilize and then treated with either rosiglitazone or troglitazone (Cayman Chemicals, Ann Arbor, MI) at two concentrations for each drug, 30× and 60× C_max_ (C_max_ is the maximum drug concentration measured in human plasma; reported C_max_ for troglitazone and rosiglitazone are 2.82 μg/mL and 0.373 μg/mL, respectively ^25^), for a total of 8 days with fresh drug added to culture medium at the 4-day medium exchange. Previously, 5-9 days of treatment with drugs was found to increase the sensitivity for drug toxicity detection without an increase in the false positive rate over 1-day drug treatment in 2D PHH / 3T3-J2 co-cultures ^21^. Both drugs were dissolved in 100% DMSO. The final DMSO concentration in the culture medium was kept at 0.1% (v/v) for both drugs, and a DMSO-only control culture was used to calculate fold changes due to the drug treatment. Albumin secretion, as a marker of hepatic dysfunction following drug treatment ^21^, was measured as described above, while overall viability of the co-culture was quantified via the PrestoBlue™ assay (Thermo Fisher) per manufacturer’s instructions. Briefly, PrestoBlue substrate was combined with culture medium at a 1:9 ratio (v:v). The mixture was added to cultures, incubated for 3 hours, and the metabolite (resazurin) was quantified via fluorescence detection (excitation/emission: 560/590 nm) on the Synergy H1 multi-mode plate reader.

### Data analysis

All findings were confirmed in 2-3 independent experiments (3-4 wells per condition and per experiment) from 2 cryopreserved PHH donors (lots HUM4055A: 54-year-old female Caucasian, and HUM4192: 16-year old female Asian; both lots purchased from Lonza, Walkersville, MD). Data processing was performed using Microsoft Excel and image analysis was performed using ImageJ ^26^. GraphPad Prism (La Jolla, CA) or Mathematica (Wolfram Research, Champaign, IL) was used for displaying results. Mean and standard deviation are displayed for all data sets. Gene expression data in microtissues were calculated as fold changes relative to gene expression data in PHHs immediately following thawing using the ΔΔC^T^ method with *GAPDH* as the housekeeping gene. Statistical significance was determined using Student’s t-test or one-way ANOVA followed by a Bonferroni pair-wise post-hoc test (p< 0.05)

## ACKNOWLEDGEMENTS

We acknowledge Regeant Panday for cell culture assistance. Funding was provided by the National Institutes of Health (1R21ES027622-01 to SRK and DKW) and the National Science Foundation (CBET-1706393 to SRK and DKW). Portions of this work were conducted in the Minnesota Nano Center, which is supported by the National Science Foundation through the National Nano Coordinated Infrastructure Network (NCCI) under award number ECCS-1542202. ALC was funded by the National Science Foundation Graduate Research Fellowship Program (00039202).

