## Supplemental Figure for "Microscale Collagen and Fibroblast Interactions Enhance Primary Human Hepatocyte Functions in 3-Dimensional Models"

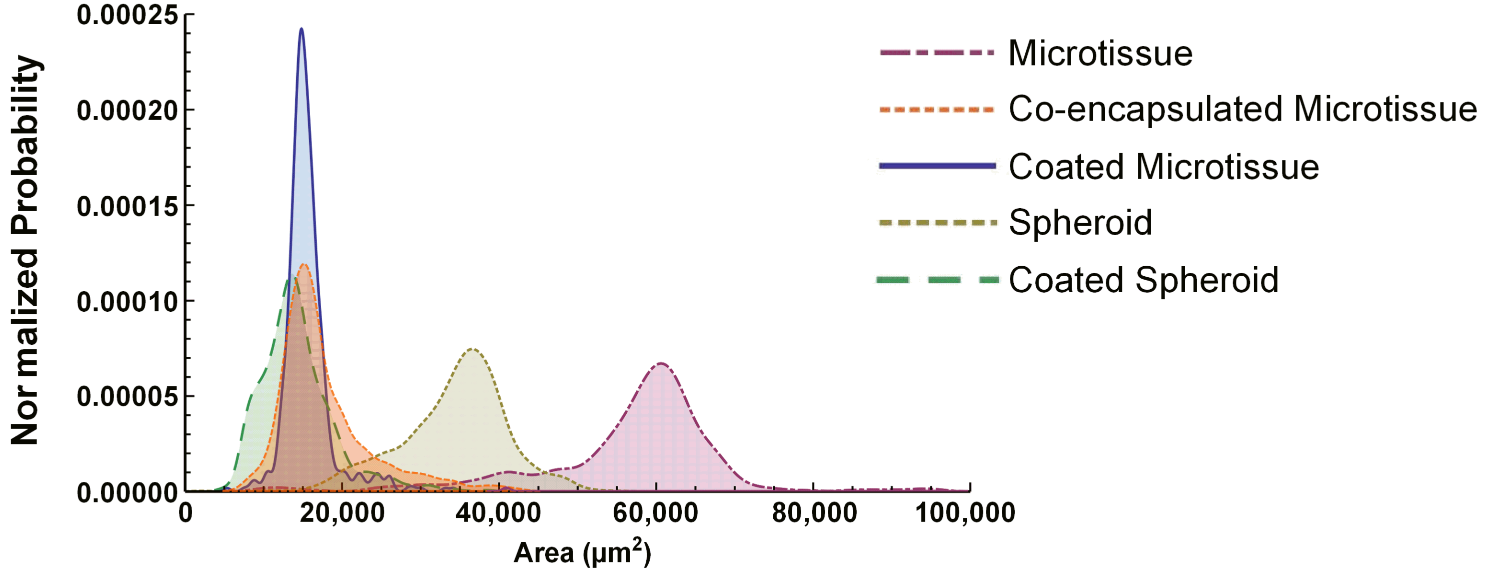


**Supplemental Figure 1. Projected surface area distributions of 3D human liver models.** Collagen-based microtissues containing encapsulated PHHs were fabricated using the droplet microfluidic device shown in Fig. 1 of the main manuscript, while self-assembled spheroids were created as shown in Fig. 2 of the main manuscript. Collagen-based microtissues containing encapsulated PHHs and 3T3-J2 fibroblasts were fabricated to create 'co-encapsulated microtissue'. 3T3-J2 fibroblasts were seeded (coated) onto the PHH microtissues or spheroids to create 'coated microtissue' and 'coated spheroid', respectively. 'Spheroid' and 'Microtissue' indicate models with only PHHs. After 21 days of culture, 375 individual constructs for each model were fixed and analyzed via phase contrast microscopy for projected surface area determinations. Shown here are the histograms of the projected surface areas for the 5 culture / co-culture models.
